# Confirmatory evidence that healthy individuals can adaptively adjust prior expectations and interoceptive precision estimates

**DOI:** 10.1101/2020.08.31.275594

**Authors:** Ryan Smith, Rayus Kuplicki, Adam Teed, Valerie Upshaw, Sahib S. Khalsa

**Affiliations:** Laureate Institute for Brain Research, Tulsa, OK, USA

**Keywords:** Interoception, Active Inference, Precision, Prior expectations, Bayesian Perception, Computational Modeling

## Abstract

Theoretical proposals have previously been put forward regarding the computational basis of interoception. Following on this, we recently reported using an active inference approach to 1) quantitatively simulate interoceptive computation, and 2) fit the model to behavior on a cardiac awareness task. In the present work, we attempted to replicate our previous results in an independent group of healthy participants. We provide evidence confirming our previous finding that healthy individuals adaptively adjust prior expectations and interoceptive sensory precision estimates based on task context. This offers further support for the utility of computational approaches to characterizing the dynamics of interoceptive processing.

## 1 Introduction

Multiple neurocomputational models of interoceptive processing have recently been put forward (e.g., (1, 2)). These models have focused largely on understanding interoception within the framework of Bayesian predictive processing models of perception. A central component of such models is the brain’s ability to update its model of the body in the face of interoceptive prediction errors (i.e., mismatches between afferent interoceptive signals from the body and prior expectations). To do so adaptively, the brain must also continuously update estimates of both its prior expectations and the reliability (precision) of afferent sensory signals arising from the body. In a recent study (3), we described a formal generative model based on the active inference framework that simulated approximate Bayesian perception within a cardiac perception (heartbeat tapping) task. We fit this model to behavioral data and found evidence that healthy individuals successfully adapted their prior expectations and sensory precision estimates during different task contexts, particularly under conditions of interoceptive perturbation. In contrast, a transdiagnostic psychiatric sample showed a more rigid pattern in which precision estimates remained stable across task conditions. As this study was the first to present such evidence, confirmatory evidence is lacking. In the present study, we attempted to replicate the previous finding in healthy participants by fitting our model to behavior on the same task in a new sample. As in our previous study, we assessed cardiac interoceptive awareness under resting conditions with different instruction sets where 1) guessing was allowed, and 2) guessing wasn’t allowed; we also 3) assessed performance during an interoceptive perturbation (inspiratory breath-hold) condition expected to increase the precision of the afferent cardiac signal (while also under the no-guessing instruction). We predicted that prior expectations for feeling a heartbeat would be reduced under the no-guessing instruction and that sensory precision estimates would increase in the breath-hold condition relative to the resting conditions. We also sought to confirm continuous relationships we previously observed between these model parameters and two facets of interoceptive awareness: self-reported heartbeat intensity (positive relationship with both parameters) and self-reported task difficulty (negative relationship with both parameters).

## 2 Methods

Data were collected from a community sample of 63 participants (47 female; mean age = 24.94, SD = 6.09) recruited via advertisements and from an existing database of participants in previous studies. Participants were screened using the Mini International Neuropsychiatric Inventory 6 or 7 (MINI) and did not meet criteria for any disorder. Our initial assessment identified some participants with poor electrocardiogram (EKG) traces, which were removed from our analyses. Final sample sizes for each condition are shown in **Table 1**.

**Table 1.**
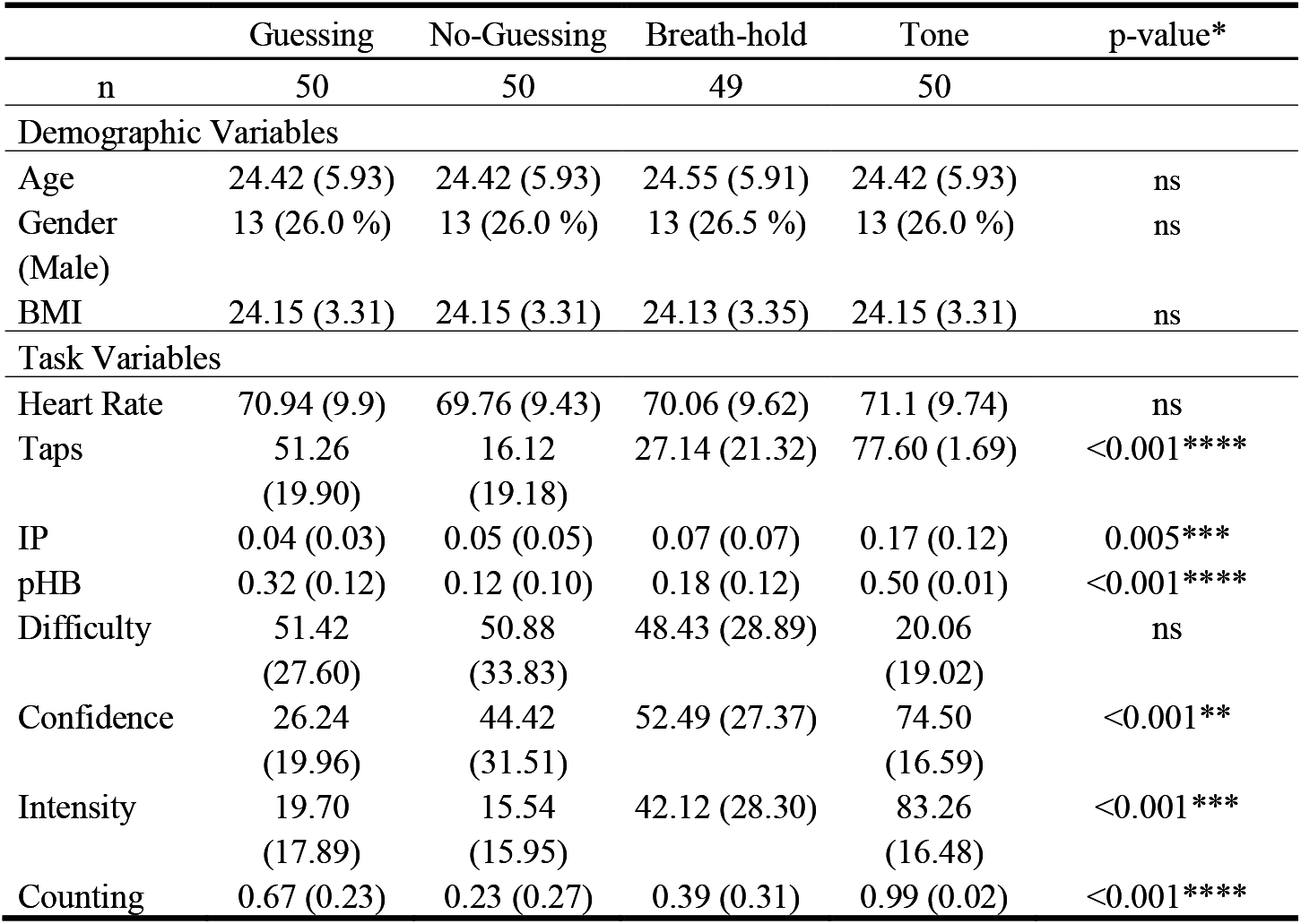
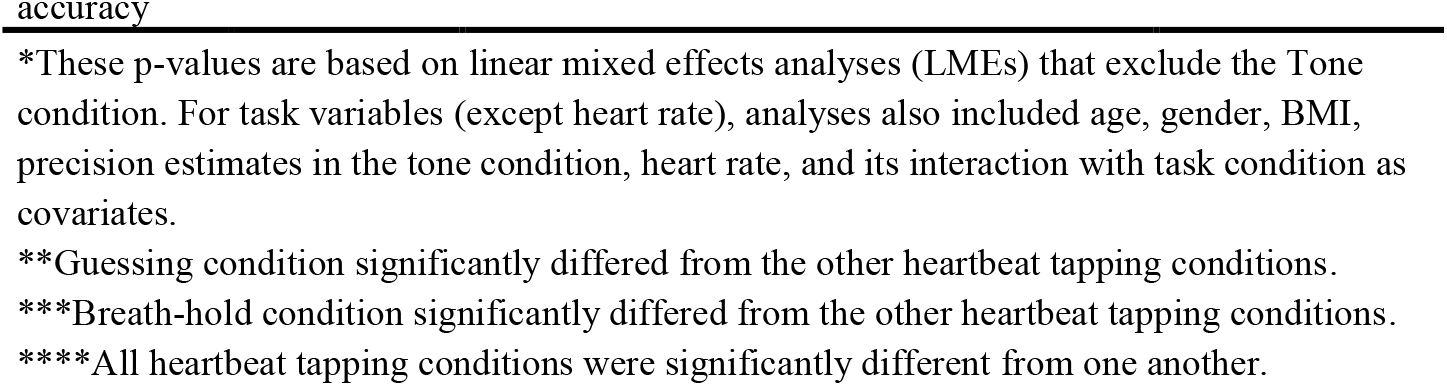
Mean and standard deviation of study variables by task condition.

Participants completed the same cardiac perception (‘heartbeat tapping’) task as in our previous study (3), wherein participants were asked to close their eyes and press down on a key each time they felt their heartbeat, and to try to mirror their heartbeat as closely as possible. Participants were not permitted to take their pulse (e.g., hold their finger to their wrist or neck) or to hold their hand against their chest. Thus, they could only base their choice to tap on their internally felt sensations. The task was repeated under multiple conditions designed to assess the influence of cognitive context and physiological perturbation on performance. In the first condition, participants were told that, even if they weren’t sure about what they felt, they should take their best guess (“guessing condition”). This condition was included because it matches a standard instruction given during heartbeat counting tasks (4). In the second condition, they were told to only press the key when they actually felt their heartbeat, and if they did not feel their heartbeat then they should not press the key (the “no-guessing” condition). In other words, unlike the first time they completed the task, they were specifically instructed not to guess if they didn't feel anything. This condition can be seen as placing an additional cognitive demand on the participant to monitor their own confidence in whether a heartbeat was actually felt; such instructions have been reported to substantially influence performance on the heartbeat counting task (5, 6). Finally, in the perturbation condition, participants were again instructed not to guess but were also asked to first empty their lungs of all air and then inhale as deeply as possible and hold it for as long as they could tolerate (up to the length of the one-minute trial) while reporting their perceived heartbeat sensations. This third condition (the “breath-hold” condition) was used in an attempt to increase the strength of the afferent cardiac signal by increasing physiological arousal. We expected 1) that cardiac perception would be poor in the guessing condition (i.e., as only roughly 35% of individuals appear to accurately perceive their heartbeats under resting conditions (7)), 2) that tapping would be more conservative in the no-guessing condition, and 3) that the breath-hold condition would result in improved performance on average (i.e., as interoceptive accuracy has been shown to increase under conditions of heightened cardiorespiratory arousal (8–10)). Directly after completing each task condition, participants were asked to rate subjective task difficulty, performance, and heartbeat intensity from 0 to 100. Participants also completed a control condition in which they tapped every time they heard a 1000Hz auditory tone presented for 100ms (78 tones, randomly jittered by +/− 10% and presented in a pattern following a sine curve with a frequency of 13 cycles/minute, mimicking the range of respiratory sinus arrhythmia during a normal breathing rate of 13 breaths per minute). This was completed between the first (guessing) and second (no-guessing) heartbeat tapping conditions. As body mass index (BMI) is a potential confound, we also measured this for each participant.

A three-lead EKG was used to assess the objective timing of participants' heartbeats throughout the task. The pulse oximeter signal was also gathered using a pulse plethysmography (PPG) device attached to the ear lobe. These signals were acquired simultaneously on a Biopac MP150 device. Response times were collected using a task implemented in PsychoPy, with data collection synchronized via a parallel port interface. EKG and response data were scored using in-house developed MATLAB code.

To model behavior, we divided each task time series into intervals corresponding to windows equally dividing the time period between each heartbeat, based on each participant’s EKG recording. Potentially perceivable heartbeats were specifically based on the timing of the peak of the EKG R-wave (signaling electrical depolarization of the atrioventricular neurons of the heart) + 200 milliseconds (ms). This 200 ms interval was considered a reasonable estimate of participants’ pulse transit time (PTT) according to previous estimates for the ear PTT (11), which signals the mechanical transmission of the systolic pressure wave to the earlobe – and was considered a lower bound on how quickly a heartbeat could be felt (and behaviorally indicated) after it occurred. We also confirmed this by measuring the PTT of each participant, defined as the distance between the peak of the EKG R-wave and the onset of the peak of the PPG waveform (usable quality median PTT values were available in 45 participants; mean = 200 ms, SD = 2 ms). The length of each heartbeat interval (i.e., the “before-beat interval” and “after-beat interval”) depended on the heart rate. For example, if two heartbeats were 1 second apart, the “after-beat interval” would include the first 500 ms after the initial beat and the “before-beat interval” would correspond to the 2nd 500 ms. The after-beat intervals were considered the time periods in which the systole (heart muscle contraction) signal was present and in which a tap should be chosen if it was felt. The before-beat intervals were treated as the time periods where the diastole (heart muscle relaxation) signal was present and in which tapping should not occur (i.e., assuming taps are chosen in response to detecting a systole; e.g., as supported by (12)). This allowed us to formulate each interval as a “trial” in which either a tap or no tap could be chosen and in which a systole or diastole signal was present (see **Fig. 1**). Each trial formally consisted of two time points. At the first time point, the model always began in a “start” state with an uninformative “start” observation. At the second time point, either a systole or diastole observation was presented, based on whether that trial corresponded to the time window before or after a systole within the participant’s EKG signal (as described above). The model then inferred the probability of the presence vs. absence of a heartbeat (corresponding to the probability of choosing whether or not to tap). At this point, the trial ended, and the next trial began with the model again beginning in the “start” state and being presented with a new systole or diastole signal, and so forth. For further details on all methods, see (3).

**Fig. 1.**
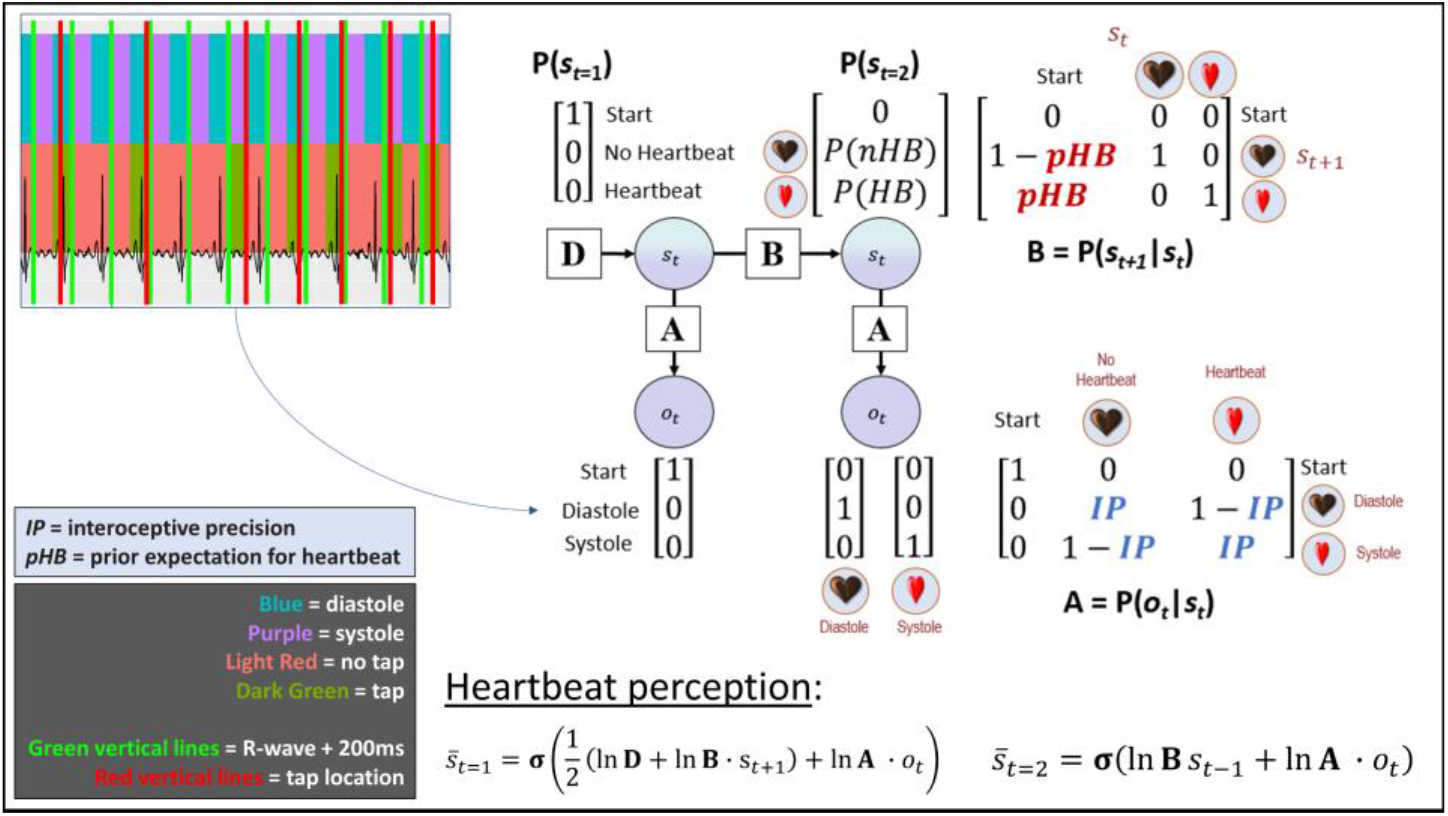
(**Upper Right**) A graphical depiction of the computational model. This is a simplified version of a commonly used active inference formulation of partially observable Markov decision processes (13), which does not explicitly model action. Systole/diastole (derived from EKG; **Upper Left**) were modeled as observations, and beliefs about the presence or absence of a heartbeat were modeled as hidden states. For simplicity, model-fitting assumed that the probability of choosing to tap corresponded to the posterior distribution over states (s – that is, the relative confidence in the presence vs. absence of a heartbeat: P(*HB*) and P(*nHB*), respectively. Estimated model parameters included: 1) interoceptive precision (*IP*) – the precision of the mapping from systole/diastole to beliefs about heartbeat/no heartbeat in the **A** matrix, which can be associated with the weight assigned to sensory prediction errors; and 2) prior expectations for the presence of a heartbeat (*pHB*). Because minimal precision corresponds to an *IP* value of .5, and both higher and lower values indicate that taps more reliably track systoles (albeit in an anticipatory or reactive manner), our ultimate measure of precision subtracted 0.5 from raw *IP* values and then took their absolute value. On each trial, beliefs about the probability of a heartbeat (corresponding to the probability of choosing to tap) relied on Bayesian inference as implemented in the “heartbeat perception” equations (**Bottom Right**). Note that, by convention in active inference, the dot product (∙) applied to matrices here indicates transposed matrix multiplication, and σ denotes a softmax (normalized exponential) function (see text for details).

To model behavior, we used a Bayesian generative model of perception (see **Fig.1**) derived from the Markov decision process (MDP) formulation of active inference (13). Unlike the full MDP model, however, we only explicitly included a generative model of perception. Observations (o) included systole, diastole, and a “start” observation (i.e., based on each individual’s EKG recording). These observations were generated by hidden (perceptual) states (s) that included either feeling one’s heartbeat or not, as well as a “start” state. The probability of choosing to tap on each trial was assumed to correspond to the posterior probability of the heartbeat state on each trial. Here, a trial formally included two timesteps: 1) a “start” time point, followed by 2) the possibility of either a systole or diastole. The matrices and equations defining the model are specified in **Fig. 1**. This model was used in conjunction with the standard SPM_MDP_VB_X routine (within the freely available SPM12 software package; Wellcome Trust Centre for Neuroimaging, London, UK, http://www.fil.ion.ucl.ac.uk/spm) to simulate cardiac response data.

The precision of the likelihood matrix **A** was controlled by an “interoceptive precision” (*IP*) parameter. Prior expectations were controlled by a parameter *pHB* within the transition matrix **B**. Note that, because each ‘trial’ was based on equally dividing the time periods before and after each heartbeat (i.e., based on each individuals EKG; resulting in alternating ‘systole’ and ‘diastole’ trials), this entails that the ‘correct’ *pHB* value would be 0.5. Both *IP* and *pHB* were estimated for each participant by finding values that maximized the likelihood of their responses using variational Laplace – that is, values that maximized the posterior probability of the heartbeat state on trials in which they chose to tap (implemented by the spm_nlsi_Newton.m parameter estimation routine available within SPM). Prior means and variances for each parameter were both set to 0.5. Because “raw” IP values (*IP_raw_*) both above and below 0.5 indicate higher precision (i.e., values approaching 0 indicate reliable anticipatory tapping, whereas values approaching 1 indicate reliable tapping after a systole), our ultimate measure of precision was recalculated by centering *IP_raw_* on 0 and taking its absolute value.

Primary confirmatory analyses included linear mixed effects analyses (LMEs) assessing the main effect of task condition on each parameter, while accounting for age, gender, BMI, heart rate, and its interaction with task condition. To help rule out the possibility that *IP* estimates are driven by differences in motor stochasticity, we also include precision estimates for the tone condition as an additional covariate. This was based on the assumption that, because the sensory signal in the tone condition is highly precise, any variability in precision estimates in the tone condition would be better explained by individual differences in random influences on behavior as opposed to perception.

## 3 Results

As in our previous study, an LME (excluding the tone condition) revealed a main effect of task condition on *IP* (F(2,95) = 5.65, *p* = .005), after accounting for age, gender, BMI, precision in the tone condition, heart rate, and its interaction with task condition (**Fig. 2**). Post-hoc Tukey comparisons indicated that *IP* was significantly greater in the breath-hold condition than in the guessing (*p* = .006) and no-guessing (*p* = .028) conditions. An identical analysis focused on *pHB* revealed the expected effect of task condition (*F*(2,95) = 56.18, *p* < .001), in which 1) *pHB* was significantly lower in the no-guessing and breath-hold conditions than in the guessing condition (*p*s < .001; note that the breath-hold condition still included the no-guessing instruction), and 2) it was higher in the breath-hold condition than in the no-guessing condition (*p* = .01).

**Fig. 2.**
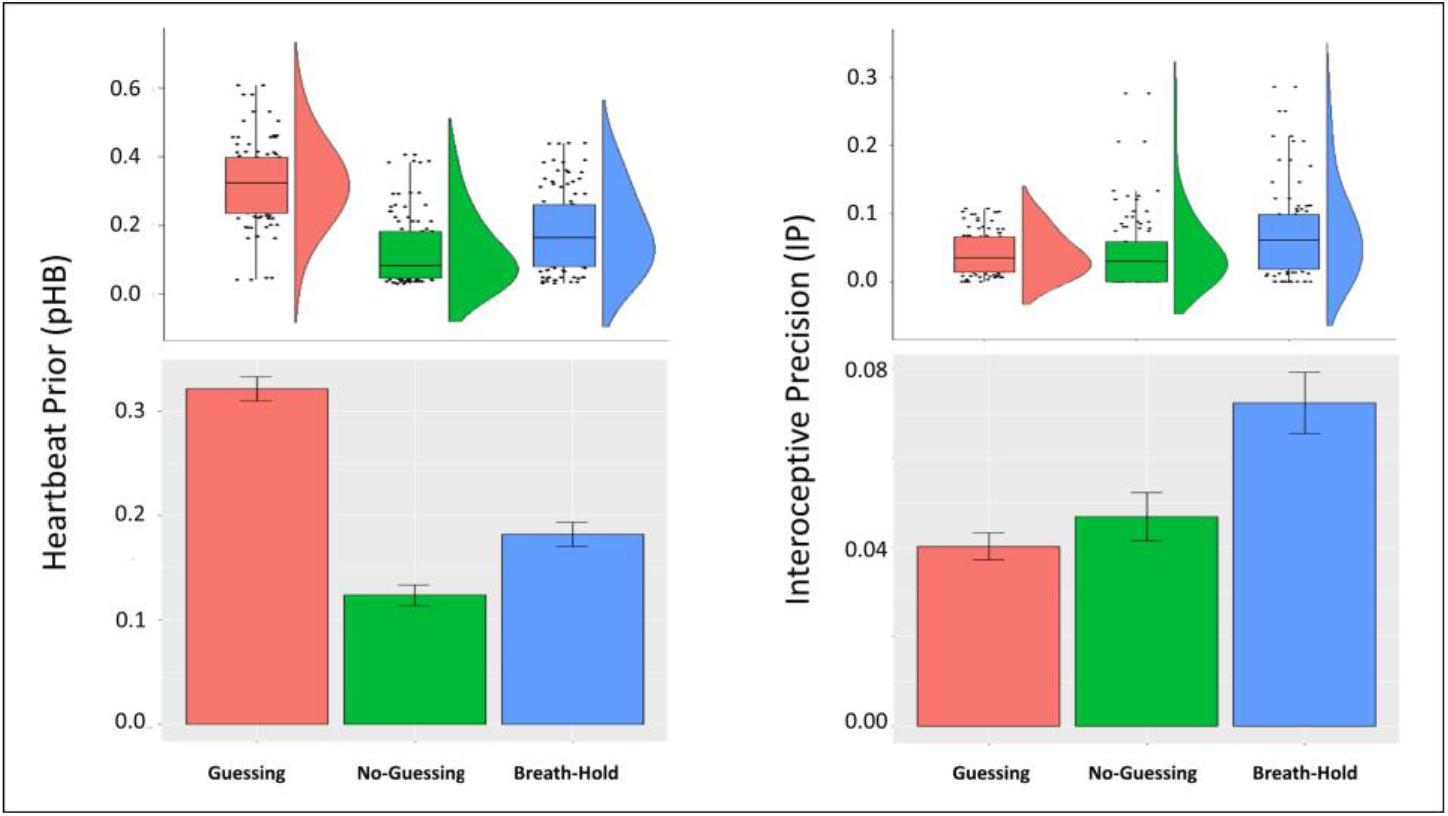
**Bottom**: Mean and standard error for prior expectations (*pHB*; **Left**) and interoceptive precision (*IP*) estimates (**Right**) by condition. **Top**: Raincloud plots depicting the same results in terms of individual datapoints, boxplots (median and upper/lower quartiles), and distributions. *pHB* was significantly lower in the no-guessing and breath-hold conditions than in the guessing condition (*p*s < .001) and it was higher in the breath-hold condition than in the no-guessing condition (*p* = .01). *IP* was significantly greater in the breath-hold condition than in the guessing (*p* = .006) and no-guessing (*p* = .028) conditions.

Secondary analyses examined the relationships between model parameters and other task variables at a threshold of *p* < .01, uncorrected (shown in **Fig. 3**). These results largely confirmed the relationships observed in our earlier study (3), including positive relationships between both *IP* and *pHB* parameters and self-reported heartbeat intensity, and negative relationships between these parameters and self-reported task difficulty. Note that expected relationships with difficulty in the breath-hold condition were not significant in this sample at our stated threshold of *p* < .01, but were significant at a more liberal threshold of *p* < .05 and had very similar correlation magnitudes as in our previous results, which were significant in that larger sample. We also confirmed that model parameters were not correlated with individual differences in median PTT.

**Fig. 3.**
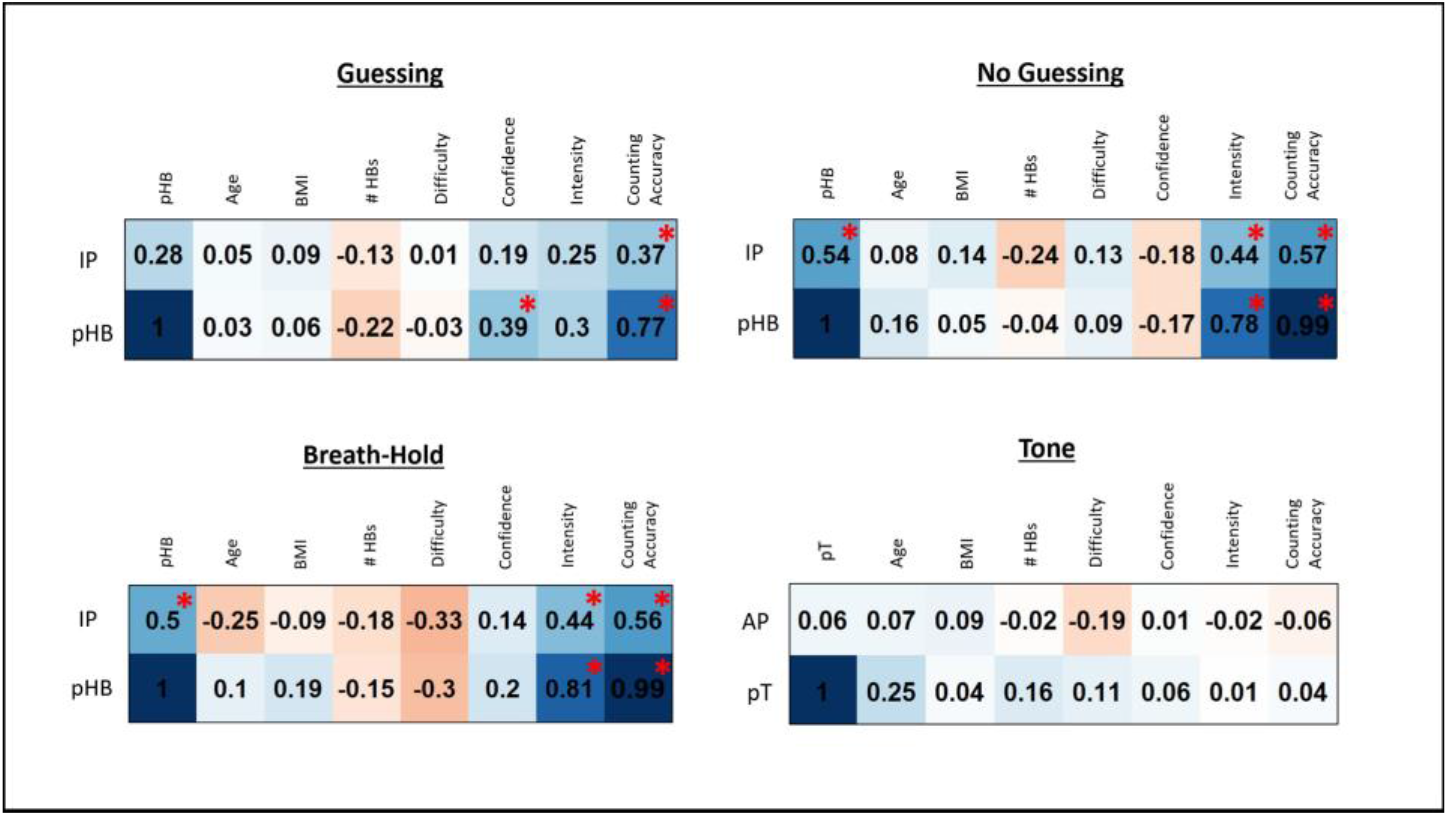
Pearson correlations between model parameters and self-report and other task-relevant variables for each task condition across all participants. *IP* = interoceptive precision parameter, *pHB* = prior expectation for heartbeat parameter, *pT* = prior expectation for tone parameter, *AP* = auditory precision, #HBs = number of heartbeats during the task condition, BMI = body mass index. For reference, correlations at *p* < .01 (uncorrected) are marked with red asterisks.

## 4 Discussion

In this study we sought, and found, confirmatory evidence that an interoceptive (inspiratory breath-hold) perturbation increased the precision estimates assigned to cardiac signals in healthy individuals. The effectiveness of the perturbation was further validated by the finding that participants reported more intense heartbeat sensations in the breath-hold condition (see **Table 1**). We further confirmed that prior expectations to feel a heartbeat were reduced when individuals were given a no-guessing instruction, and that both parameters correlated with self-report measures in predicted directions. This replication represents an important step towards empirically advancing our understanding of the computational dynamics underlying interoception. Future work remains to confirm our other previous finding – that interoceptive precision is *not* adjusted across conditions within psychiatric disorders (3). If this latter result is replicated in future work, it would support the use of our novel interoceptive modelling and modelfitting approach as an important new avenue for computationally phenotyping patient populations at the individual level.

## References

1. Barrett L, Simmons W. Interoceptive predictions in the brain. Nature reviews Neuroscience. 2015;16:419–29.

2. Smith R, Thayer JF, Khalsa SS, Lane RD. The hierarchical basis of neurovisceral integration. Neurosci Biobehav Rev. 2017;75:274–96.

3. Smith R, Kuplicki R, Feinstein J, Forthman KL, Stewart JL, Paulus MP, et al. A Bayesian computational model reveals a failure to adapt interoceptive precision estimates across depression, anxiety, eating, and substance use disorders. medRxiv. 2020:2020.06.03.20121343.

4. Pollatos O, Herbert BM, Matthias E, Schandry R. Heart rate response after emotional picture presentation is modulated by interoceptive awareness. Int J Psychophysiol. 2007;63(1):117–24.

5. Desmedt O, Luminet O, Corneille O. The heartbeat counting task largely involves non-interoceptive processes: Evidence from both the original and an adapted counting task. Biol Psychol. 2018;138:185–8.

6. DeVille DC, Kuplicki R, Stewart JL, Tulsa I, Aupperle RL, Bodurka J, et al. Diminished responses to bodily threat and blunted interoception in suicide attempters. Elife. 2020;9.

7. Khalsa SS, Lapidus RC. Can Interoception Improve the Pragmatic Search for Biomarkers in Psychiatry? Front Psychiatry. 2016;7:121.

8. Khalsa SS, Rudrauf D, Sandesara C, Olshansky B, Tranel D. Bolus isoproterenol infusions provide a reliable method for assessing interoceptive awareness. Int J Psychophysiol. 2009;72(1):34–45.

9. Hassanpour MS, Yan L, Wang DJ, Lapidus RC, Arevian AC, Simmons WK, et al. How the heart speaks to the brain: neural activity during cardiorespiratory interoceptive stimulation. Philos Trans R Soc Lond B Biol Sci. 2016;371(1708).

10. Schandry R, Bestler M, Montoya P. On the relation between cardiodynamics and heartbeat perception. Psychophysiology. 1993;30(5):467–74.

11. Allen J, Murray A. Age-related changes in the characteristics of the photoplethysmographic pulse shape at various body sites. Physiol Meas. 2003;24(2):297–307.

12. Ring C, Brener J. The temporal locations of heartbeat sensations. Psychophysiology. 1992;29(5):535–45.

13. Da Costa L, Parr T, Sajid N, Veselic S, Neacsu V, Friston K. ACTIVE INFERENCE ON DISCRETE STATE-SPACES – A SYNTHESIS. arXiv. 2020:2001.07203v2 [q-bio.NC]

